# The genetic variation of lactase persistence alleles in northeast Africa

**DOI:** 10.1101/2020.04.23.057356

**Authors:** Nina Hollfelder, Hiba Babiker, Lena Granehäll, Carina M Schlebusch, Mattias Jakobsson

## Abstract

Lactase persistence (LP) is a well-studied example of a Mendelian trait under selection in some human groups due to gene-culture co-evolution. We investigated the frequencies of genetic variants linked to LP in Sudanese and South Sudanese populations. These populations have diverse subsistence patterns, and some are dependent on milk to various extents, not only from cows, but also from other livestock such as camels and goats. We sequenced a 316bp region involved in regulating the expression of the *LCT* gene on chromosome 2, which encompasses five polymorphisms that have been associated with LP. Pastoralist populations showed a higher frequency of LP-associated alleles compared to non-pastoralist groups, hinting at positive selection also in northeast African pastoralists. There was no incidence of the East African LP allele (−14010:C) in the Sudanese groups, and only one heterozygote individual for the European LP allele (−13910:T), suggesting limited recent admixture from these geographic regions. Among the LP variants, the −14009:G variant occurs at the highest frequency among the investigated populations, followed by the −13915:G variant, which is likely of Middle Eastern origin, consistent with Middle Eastern gene-flow to the Sudanese populations. The Beja population of the Beni Amer show three different LP-variants at substantial and similar levels, resulting in one of the greatest frequencies of LP-variants among all populations across the world.

## Introduction

Lactase persistence (LP) is the ability to digest the milk sugar, lactose, at an adult age. The phenotype is associated with several single nucleotide polymorphisms (SNPs) that are located 13.9kb upstream of the lactase gene (*LCT*) in an associated enhancer element. Currently, we know of at least five variants that are clearly associated with the LP phenotype (Ségurel and Bon, 2017). The best known case is the −13910:C*>*T polymorphism (rs4988235), which is strongly associated with LP in populations of European ancestry (Enattah *et al*., 2002) and has been under strong recent selection, likely co-evolving with dairy farming (Bersaglieri *et al*., 2004).

The LP phenotype has been found at greater frequencies in milk-drinking pastoralist populations than non-pastoralist populations (Gerbault *et al*., 2011; Holden and Mace, 1997; Itan *et al*., 2010), and is also common in pastoralist African societies (Tishkoff *et al*., 2007). However, LP occurs in populations that do not carry the derived −13910:T allele, specifically in the Middle East and Eastern Africa, therefore, the thoroughly investigated −13910:C*>*T polymorphism is not the causal variant in these populations (Myles *et al*., 2005). Other SNPs have been identified to be the putative causal variants in these regions: −13907:C*>*G (rs41525747) in Ethiopia and Saudi- Arabia, −13915:T*>*G (rs41380347) in Saudi- Arabia, −14009:T*>*G (rs869051967) in African Arab groups, and −14010:G*>*C (rs145946881) in Kenya and Tanzania (Ingram *et al*., 2009; Jones *et al*., 2013; Liebert *et al*., 2016; Priehodová *et al*., 2014; Ranciaro *et al*., 2014; Tishkoff *et al*., 2007). These polymorphisms have been shown to increase *LCT* promoter expression in vitro (Enattah *et al*., 2008; Jensen *et al*., 2011; Jones *et al*., 2013; Liebert *et al*., 2016; Olds *et al*., 2011; Tishkoff *et al*., 2007), and the −13910:C*>*T variant was recently identified as the putative causal variant for LP in a GWAS study in the Fulani population of the African Sahel/Savannah belt (Vicente *et al*., 2019). There is evidence for a selective sweep on −14010:G*>*C (Tishkoff *et al*., 2007) that shows a stronger selection coefficient in the Massai (MKK) than the allele −13910:T shows in the European (CEU) population (Altshuler *et al*., 2010; Schlebusch *et al*., 2013), pointing to a strong increase in fitness for LP individuals in African pastoralist populations.

LP-associated SNPs have been reported in Northeast Africa (Enattah *et al*., 2008; Hassan *et al*., 2016; Tishkoff *et al*., 2007) and there is linguistic and archaeological evidence that cow-herding has been practiced in northeast Africa for at least four thousand years (Ehret, 1979; Smith, 1992). The development of farming in northeast Africa depended on the climatic conditions. While the wetter conditions along the Nile allowed for crop farming and settlement, pastoralism with a semi-nomadic lifestyle was developed in the drier Savannah/Sahel regions (Haaland and Haaland, 2013). The pastoralist Beja populations of Sudan have been shown to have a high prevalence of LP (Bayoumi *et al*., 1981; Tishkoff *et al*., 2007) and moderately high allele frequencies of LP-associated alleles compared to neighboring populations, which could have arisen due to a selection event (Ranciaro *et al*., 2014). The Nilotic populations of current-day South Sudan are dairy consuming pastoralists, which have been shown to be lactase persistent in low frequencies (Bayoumi *et al*., 1981, 1982; Tishkoff *et al*., 2007), but no alleles associated with LP have this far been found (Hassan et al., 2016; Tishkoff et al., 2007).

To deepen our understanding of LP in Northeast Africa and the associated variants, we sequenced a 316bp region spanning all known SNPs associated with LP, in 221 individuals from 18 Sudanese and South Sudanese (SASS) populations. Combining this data with previously published high-density genome-wide genotyping data of the same individuals (Hollfelder et al., 2017), we were able to investigate the allele frequencies of the LP associated SNPs, their haplotype backgrounds and to scan for signals of selection.

## Results and Discussion

### Allele frequencies

In total, we identified ten different polymorphisms in this study (Table 1). We detected four of the five LP-associated alleles (−13907:G, −13910:T, −13915:G, and −14009:G) and their frequencies per population are shown in Table 2. None of the LP-associated SNPs are significantly deviating from Hardy-Weinberg equilibrium in the investigated SASS populations.

**Table 1.**
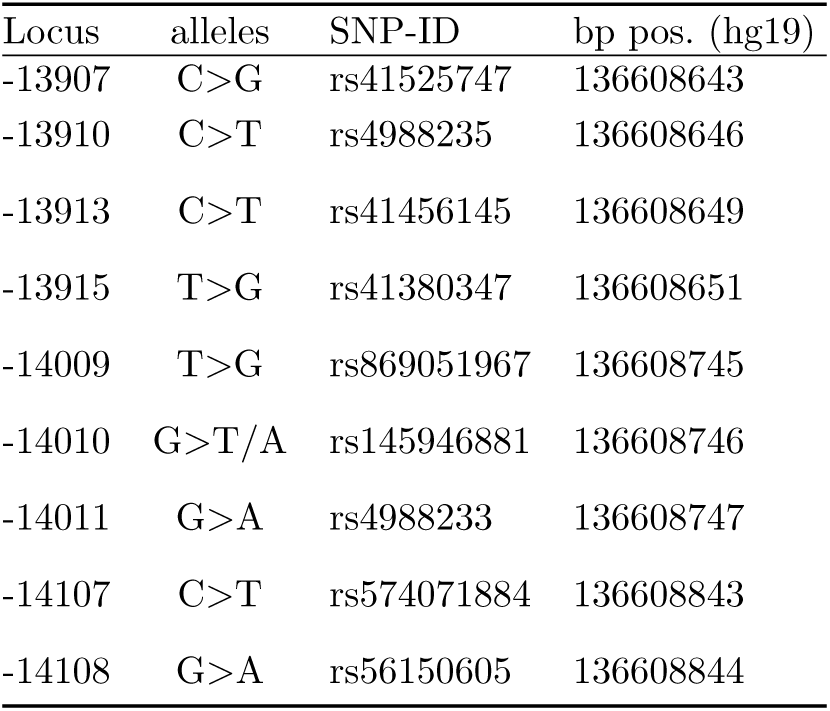
SNPs identified on the targeted sequences

**Table 2.**
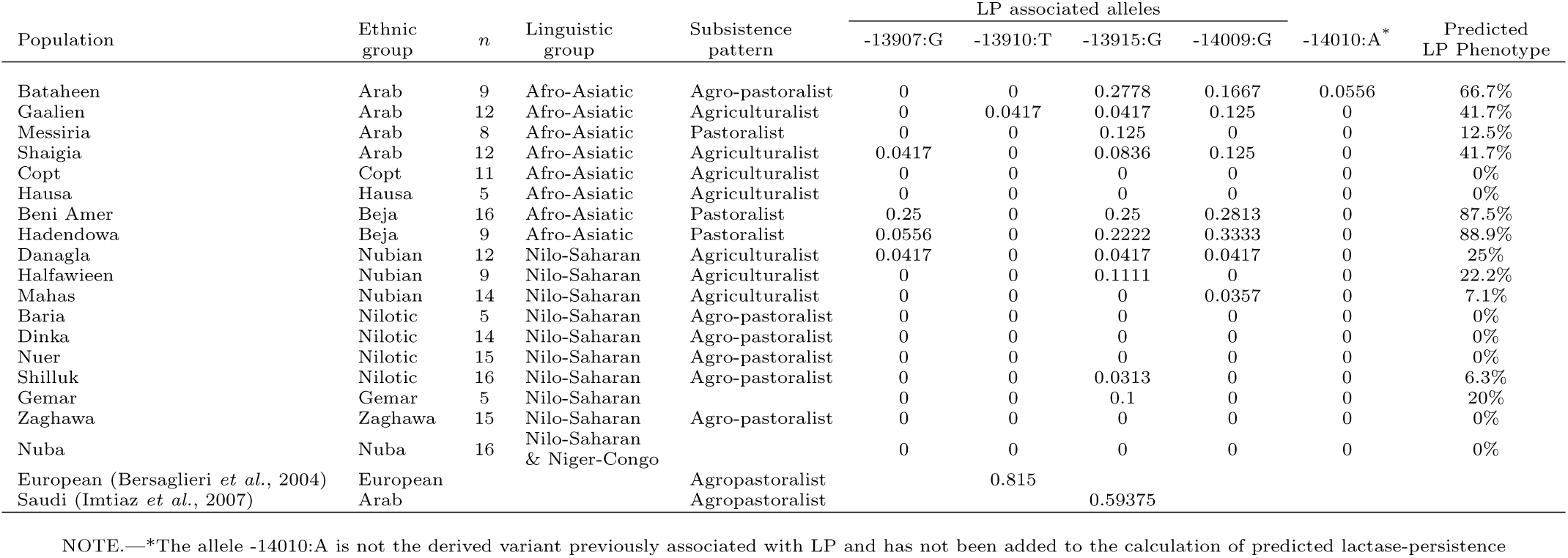
:Population information and allele frequencies of alleles at positions associated with LP.

We found a Bataheen individual with a derived adenine allele at position −14010 (5.6% allele frequency in the population, Table 2). While this is not a novel allele, it occurs at very low frequencies and has not been associated with LP. The LP-associated allele −14010:C has been shown to occur at its highest frequency in the Afro-Asiatic and Nilosaharan pastoralist populations of East Africa (Schlebusch *et al*., 2013; Tishkoff *et al*., 2007; Wagh *et al*., 2012), and was also found in southern Africa, where it was introduced through gene flow from East Africa (Breton *et al*., 2014; Coelho *et al*., 2009; Macholdt *et al*., 2014; Torniainen *et al*., 2009).

The allele associated with LP in Europeans, −13910:T, was almost completely absent from the investigated populations, except for one heterozygote Gaalien individual (Table 2). The −13910:T allele has previously been detected in African populations, as a result of European gene flow, and has also been reported to occur in populations of Sudan in low frequencies (∼0.01 allele frequency) such as the Shokrya (Hassan *et al*., 2016), the Gaalien (Ingram *et al*., 2007), and the Beni Amer (Jones *et al*., 2015), as well as in higher frequency in the Fulani population that spread across the Sahel/Savannah belt (0.23– 0.48 allele frequency, Enattah *et al*., 2007; Hassan *et al*., 2016; Ingram *et al*., 2007; Lokki *et al*., 2011; Ranciaro *et al*., 2014; Vicente *et al*., 2019).

The LP-associated alleles −13907:G, −13915:G, and −14009:G appear in frequencies up to 0.34 in SASS, mainly in Arab, Nubian and Beja populations of Sudan (Table 2). The LP-associated allele −13915:G has previously been found in the Middle East, where it likely originated (Priehodová *et al*., 2017), and in East Africa (Enattah *et al*., 2008; Imtiaz *et al*., 2007; Ingram *et al*., 2007, 2009; Priehodová *et al*., 2017; Tishkoff *et al*., 2007). It has been shown that −13915:G appears at higher frequencies in nomadic populations compared to sedentary populations (Priehodová *et al*., 2017). The allele likely spread from the Middle East to Africa through the nomadic Bedouin population, where it occurs at high frequencies (Ingram *et al*., 2007; Priehodová *et al*., 2014). The derived allele of −13915 is found in all Arab populations of Sudan (4.2- 27.8%, Table 2). The observed allele frequencies are similar to previously reported values for the Gaalien and Shaigia (Enattah *et al*., 2008; Hassan *et al*., 2016; Ingram *et al*., 2007). This is the only LP-variant present in the Messiria population (12.5%), an Arab population from southwest Sudan. It is also found in the Beja and Nubian populations, concurrent with previous results (Hassan *et al*., 2016). We furthermore find −13915:G at 3.1% in the Shilluk and at 10% in the Gemar. The allele frequency of −13915:G correlates significantly (*ρ*=0.5880782, p=0.01026) with the Middle Eastern admixture proportions of the carrier populations (Figure 2).

The derived allele of −13907 has been found in populations of the Sudan and East Africa (Ingram *et al*., 2007; Jones *et al*., 2013; Ranciaro *et al*., 2014; Tishkoff *et al*., 2007). The Beja populations have been shown to carry the highest frequencies of this allele, along with Afro-Asiatic Kenyans (Tishkoff *et al*., 2007). The −13907:G allele has also been observed in low frequency in sedentary Arab populations of Sudan (Enattah *et al*., 2008; Ingram *et al*., 2007; Ranciaro *et al*., 2014) and in the Danagla (*<*10%) (Ingram *et al*., 2007). We observed this allele only at low frequency (*<*5%) in the Shaigia Arab population and the Nubian Danagla. A previous study of LP in Sudan (Hassan *et al*., 2016), also found only low frequencies (*<*5%) of this allele in populations other than the Beja, where we observed allele frequencies up to 25%.

The LP-associated polymorphism −14009:G has its highest reported occurrence in the Beja populations of Sudan, but is also found in smaller frequencies in other populations such as African Arab groups and the populations of the Middle East and East Africa (Ingram *et al*., 2009; Jones *et al*., 2013, 2015; Liebert *et al*., 2016; Priehodová *et al*., 2014; Ranciaro *et al*., 2014). In this study, the highest frequency of this allele is also found in the Beja populations, but it is also the most common occurring LP-associated allele found in SASS populations.

The Beni Amer of the Beja show the highest frequencies of LP-associated alleles among the tested populations (Figure 1). Both −13907:G and −13915:G appear at 25%, and −14009:G has a frequency of 28.1%. The Hadendowa have a higher frequency of the derived allele −14009:G (33.3%), but lower frequency of −13915:G (22.2%) and −13907:G (5.6%) than the Beni Amer. The −13907:G variant was previously reported at even higher frequencies in the Hadendowa than observed here (Ranciaro *et al*., 2014). The comparatively high allele frequencies of LP-associated alleles leads to the highest prediction of LP-phenotype of close to 90% in the Beja populations (Table 2). Previous studies have registered the LP-phenotype to be 64%-100% in the Beni Amer and 82% in the Hadendowa (Bayoumi *et al*., 1981; Holden and Mace, 1997; Tishkoff *et al*., 2007).

**FIG. 1.**
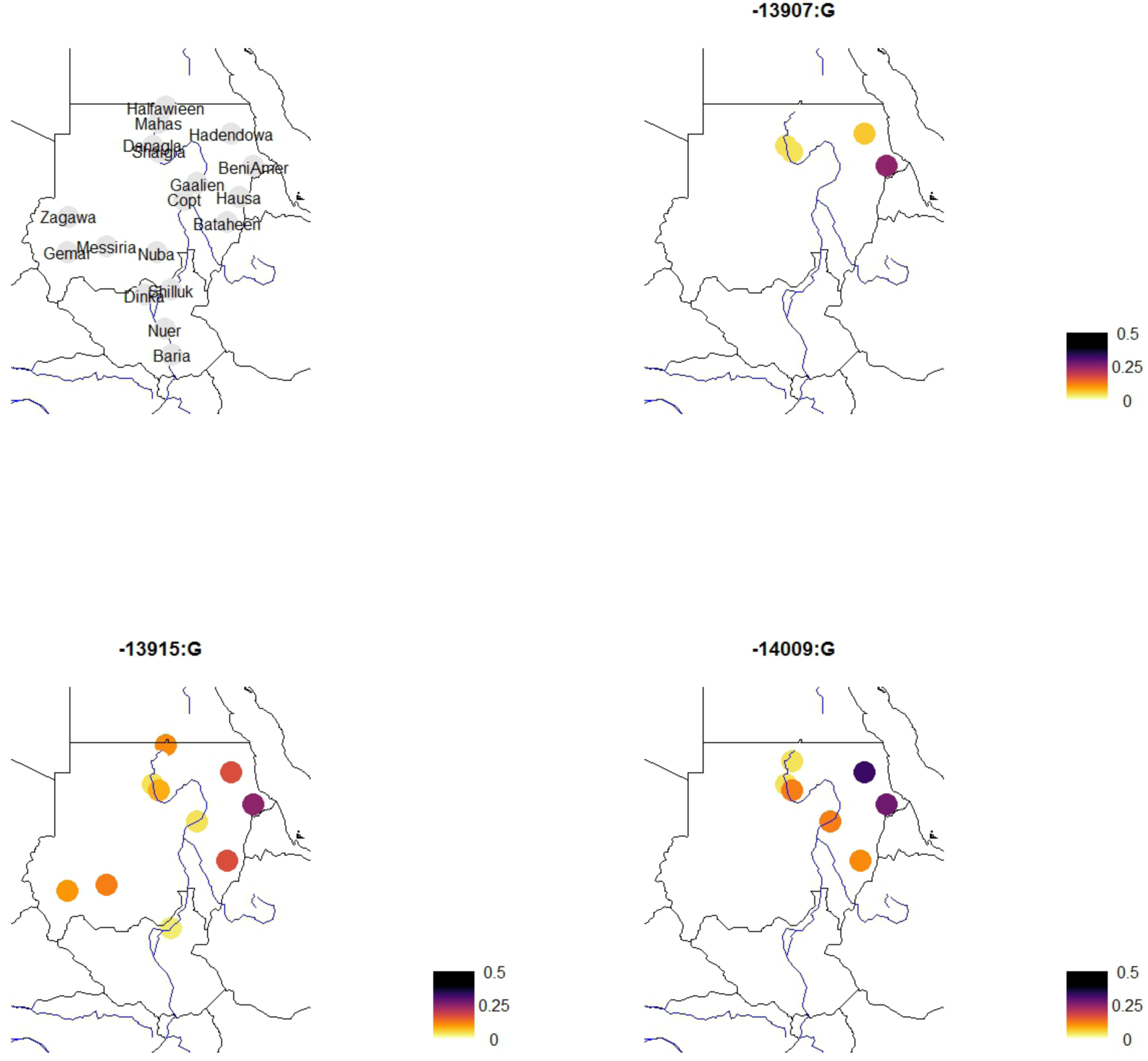
Allele frequency distribution of the three LP-associated alleles found in multiple SASS populations. A distribution of these alleles (including −13010:T and −14010:C) in Africa can be seen in figure S6.

**FIG. 2.**
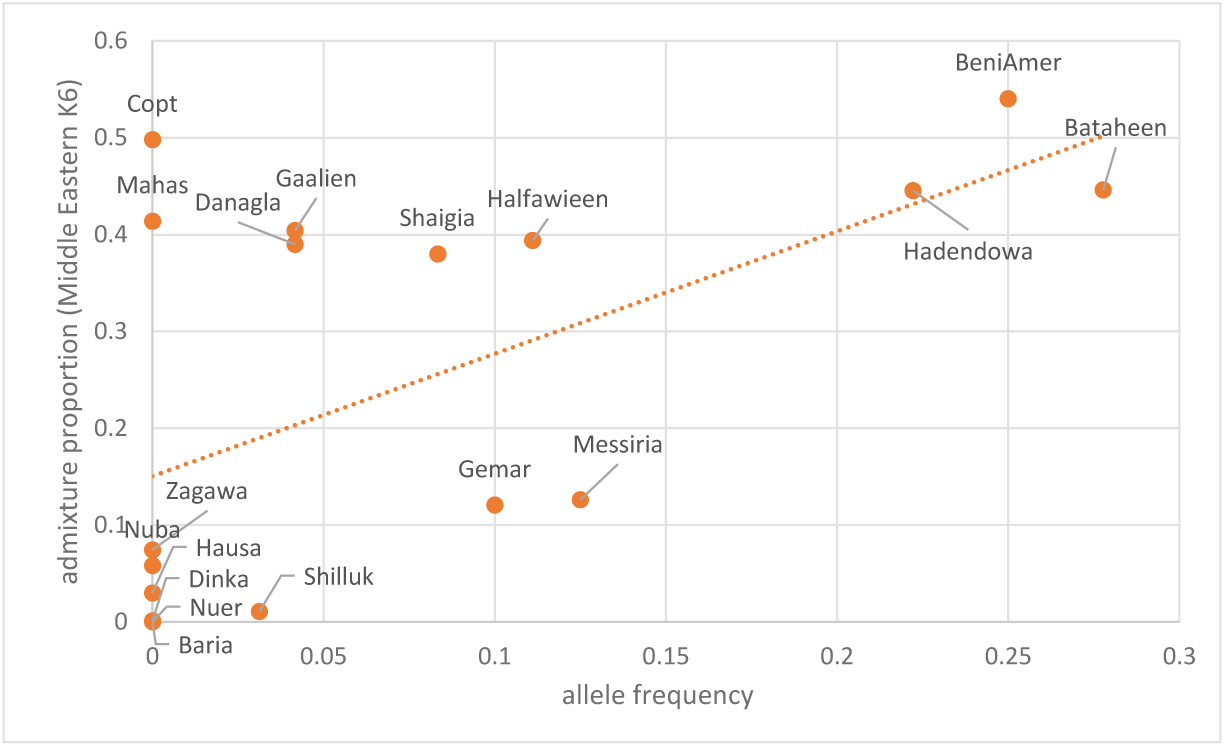
Comparison of allele frequency of −13915:G to the Middle Eastern ancestry component assuming 6 clusters in an ADMIXTURE analyses with worldwide populations (Hollfelder *et al*., 2017, Fig. S3).

The genetic differentiation, that has been identified previously between the Arabs of central/north Sudan and the Messiria (Babiker *et al*., 2011; Hollfelder *et al*., 2017), is also seen in the frequency of the LP alleles. The derived allele for −14009 is found in the Bataheen, Gaalien and Shaigia at 0.125-0.167 frequency but not in the Messiria (site specific 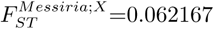 to 0.113881, where X is one of the other Sudanese Arab populations). Both groups have previously been shown to be genetically close to surrounding populations and resulted through an admixture process between local populations and migrating Middle Eastern populations leading to the emergence of an Arab ethnic affiliation (Hollfelder *et al*., 2017). The Messiria are part of the Baggara Arabs, a collective term for nomadic tribes of Kordofan that are dairy farming pastoralists (Bayoumi *et al*., 1981). Priehodová *et al*. (2017) hypothesized that there were two directions of Middle Eastern gene flow, one entered along the Nile giving rise to the Arab populations that reside along the Nile, while the other came from Lake Chad, forming the Baggara Arabs. This could potentially explain the absence of the other LP-associated alleles, other than −13915:G, in the Messiria. Alternatively, through the lower levels of admixture seen in the Messiria, only −13915:G might have persisted.

The Nubians show low frequencies of the LP-associated alleles. The Danagla have three individuals with one heterozygote derived LP- allele each (4.2% frequency of each −13907:G, −13915:G, and −14009:G). A previous study has not observed −13915:G in the Danagla (Ingram *et al*., 2007), but −13907:G was observed at 8% (Ingram *et al*., 2007). The Halfawieen only carry derived alleles of −13915 (11.1%), concurrent with previous results (Hassan *et al*., 2016), and the Mahas have one individual with heterozygote state of the −14009 polymorphism (3.6%). The −13915:G allele was previously observed at 0.038- 0.167 allele frequency in the Mahas (Enattah *et al*., 2008; Hassan *et al*., 2016), but no derived allele was found in the Mahas in this study. The predicted frequency of lactose digesters (Table 2) is in agreement with frequencies found in a study identifying lactose digesters through the hydrogen breath test (Bayoumi *et al*., 1981). The Nubians and Sudanese Arab populations have similar levels of Middle Eastern admixture, however, the Nubians show lower frequencies of the LP-associated alleles. The genetic differentiation of the LP-associated alleles between Nubians and central Sudanese Arabs is higher than 0.05 in three of the nine pairwise comparisons, both pairing a Nubian with the Bataheen population. The Bataheen also show differentiation in the LP-associated alleles to the Gaalien (*F*_*ST*_ *>* 0.05). The Bataheen show the highest frequencies of LP-associated alleles and have the highest predicted LP phenotype of the Nubian and Arab populations. Assuming that the non-African admixture into all Arab and Nubian population occurred in the same event, it is possible that there is a stronger selective pressure on the LP-associated region in the camel-breeding Bataheen than the other populations, that are more reliant on agriculture (Bayoumi *et al*., 1981; Hassan *et al*., 2016).

No LP-associated alleles were found in the Nilotic populations of South Sudan. Due to the close proximity of SASS populations to East Africa it is surprising that there is no evidence of the derived −14010:C allele in the SASS populations. This absence in Nilotic South Sudanese populations, despite the occurrence in Nilotic Tanzanians and Kenyans, where it is significantly associated with LP, has previously been noted (Tishkoff *et al*., 2007), and is in agreement with a previous study, that found the South Sudanese Nilotes to have remained largely isolated (Hollfelder *et al*., 2017). The lack of LP-associated alleles in the agro-pastoralist Nilotic populations has been observed before (Hassan *et al*., 2016; Tishkoff *et al*., 2007) despite the intermediate prevalence of lactose-digesters (*>*20%) in tested Nilotic populations (Bayoumi *et al*., 1981, 1982; Tishkoff *et al*., 2007). This might be indicative of unknown LP associated variants in the Nilotic populations. LP associated alleles are also absent in the Hausa of Sudan, although a Hausa population of Cameroon had previously shown 0.139 allele frequency of −13910:T (Mulcare *et al*., 2004). In an early study of lactose digesters in Sudan (Bayoumi *et al*., 1981) the Nuba and the Messiria also showed higher LP-phenotypes than predicted in this study. These populations are genetically close to the Nilotic populations (Hollfelder *et al*., 2017) and LP might be driven by the same unknown mechanism/mutations as in the Nilotes.

Additional SNPs were found within the 316 bp region that have not been associated with LP (Table 1). The −13913:C*>*T (rs41456145) polymorphism was found in heterozygote state in one Mahas and one Copt individual (allele frequencies: 0.0357 and 0.0454). Although this SNP is inside the Oct-1 binding site (Ingram *et al*., 2007), it does not appear to have an effect on LP (Jones *et al*., 2013). This SNP has previously been found in the Gaalien of Sudan and Fulani of Cameroon (Ingram *et al*., 2007), Khoe-San populations at frequencies up to 0.075 (Breton *et al*., 2014; Macholdt *et al*., 2014), and Ethiopian populations up to 0.09 (Jones *et al*., 2013). One Bataheen individual was found to be heterozygote for −14011:G*>*A (rs4988233) (0.0556). This SNP has been shown to influence promoter activity in vitro (Liebert *et al*., 2016) and has previously been observed in European and Middle Eastern populations (Lember *et al*., 2006; Liebert *et al*., 2016), Bantu-speaking populations of southern Africa (Macholdt *et al*., 2014), as well as Ethiopian populations (Jones *et al*., 2013). The allele −14107:T (rs574071884) was found in one instance in a Beni Amer individual (allele frequency: 0.03125). This SNP has previously been found in Xhosa and Ghana populations (Torniainen *et al*., 2009), the Fulani of Mali (Lokki *et al*., 2011), Shuwa Arabs of Chad (Priehodová *et al*., 2014), and Bantu populations of Southern Africa (Macholdt *et al*., 2014). One allele of −14108:A (rs56150605) has been found in the Danagla. This allele has previously been encountered in the Gaalien of Sudan in very low frequency (Enattah *et al*., 2008).

### Haplotype Structure

We created a plot showing the allelic state of each SNP in the populations containing the three LP-associated alleles found in moderate frequencies in the investigated populations: −13907, −13915, and −14009 (Figure 5). As observed before (Tishkoff *et al*., 2007), the LP-associated SNPs are found in distinct haplotype blocks, and have therefore evolved independently. Furthermore, bifurcation plots visualize the extension of the haplotypes surrounding the LP-associated alleles in the Beja population, who carry the highest number of LP-associated alleles (Figure 3). These plots might over-represent haplotypes due to the allelic drop-out, as might be the cause for a particular long run of homozygosity around the position of the LP alleles in a Hadendowa individual, who is homozygous for −14009:G (Figures 3 and S10).

**FIG. 3.**
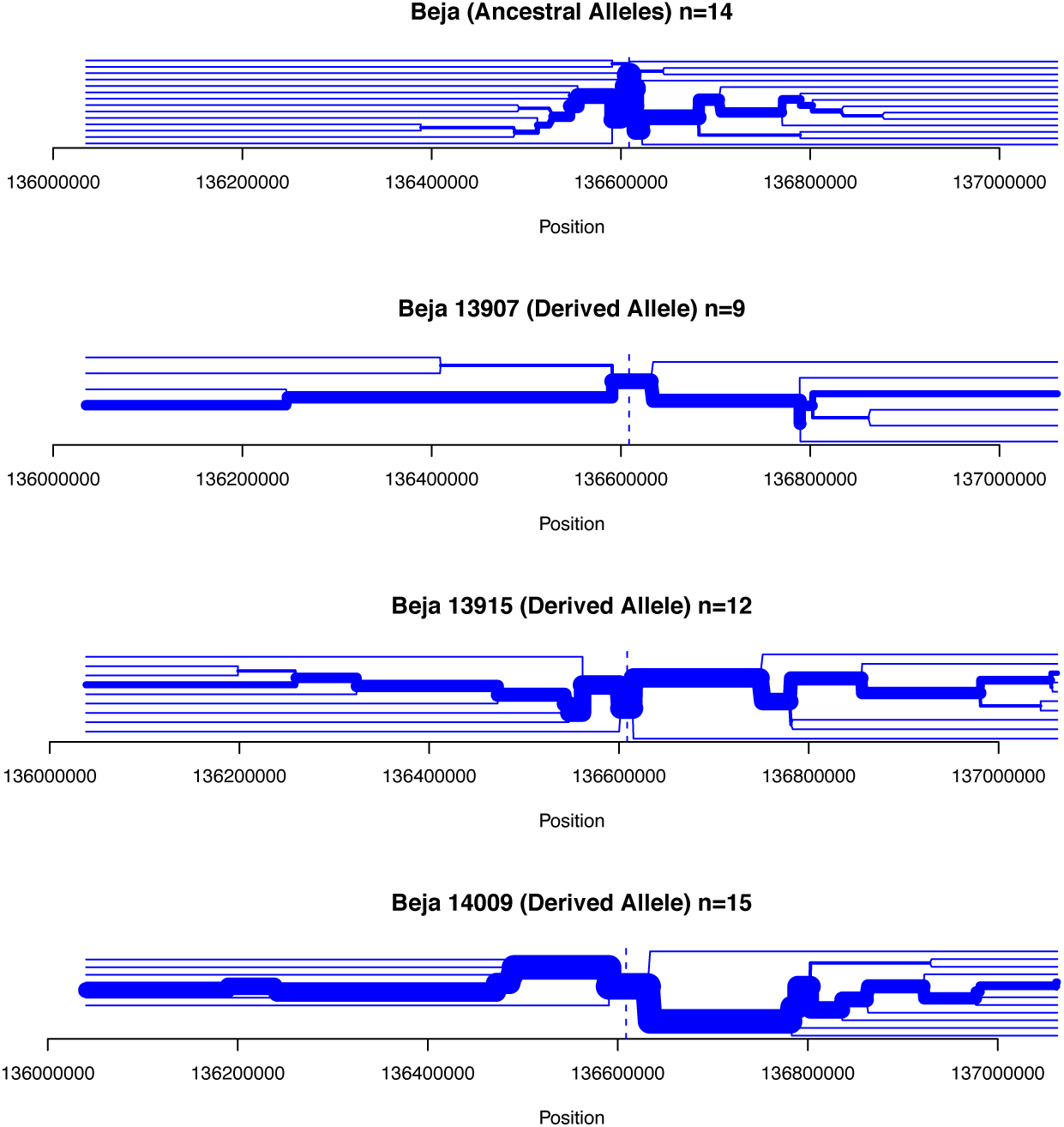
Bifurcation plots for the Beja populations. The topmost plot is centered around position −13907 and contains only haplotypes that have non of the derived LP-associated alleles.

### Selection Scan

We computed the LSBL-statistic (Shriver *et al*., 2004) across chromosome 2 and for each SASS population, as well as the MKK and CEU population from the 1000 Genomes Project dataset (1000 Genomes Project Consortium, 2015) (Figures 4, S1, S2, S3, S4, and S5), to search for signals of positive selection. In both datasets, the area around the LP-associated polymorphisms is a clear outlier in MKK and CEU, which have previously been shown to be subjected to strong positive selection (Bersaglieri *et al*., 2004; Schlebusch *et al*., 2013). Both Beja populations show increased LSBL-signals in one of the neighboring windows. It is likely that selective event is responsible for the high frequency of lactose digesters. The increase of the frequency of associated alleles surrounding the target of selection is consistent with a selective sweep. We might be observing three independent sweeps affecting the three LP-associated alleles in the same region, which appears similar to a soft sweep and is difficult to detect. This sweep might, however, have differentiated the regions surrounding the LP-associated alleles enough to increase the LSBL above the threshold. Two other regions on chromosome 2 are distinguished from the comparative populations and affect more than four populations (Table S1).

## Conclusion

LP-associated persistence alleles from Europe (−13910:T) and East Africa (−14010:C) have been used to track migration patterns of African populations (Ben Halima *et al*., 2017; Breton *et al*., 2014; Enattah *et al*., 2007; Myles *et al*., 2005). Sudanese populations have been shown to be recipients of non-African gene flow, likely from a Middle Eastern source (Hollfelder *et al*., 2017). The absence of the European and East African LP-allele (−13910:T and −14010:C) suggests negligible amounts of recent gene-flow from these regions into the populations of Sudan and South Sudan, while the occurrence of the allele associated with LP in the Middle East (−13915:G) is consistent with recent gene-flow from the Middle East into Sudan.

Even though this study investigated a range of Nilotic populations, no LP-associated SNPs were detected in these agropastoralist populations, that have been shown to be able to digest milk in hydrogen breath tests (Bayoumi *et al*., 1981, 1982; Tishkoff *et al*., 2007). Further studies on Nilotic populations will reveal more about the substructure in northeast Africa (Hollfelder *et al*., 2017) and can be informative about the underlying biology of LP in Nilotic populations. The traditionally pastoralist Beja people have been shown to have among the highest number of lactose digesters in the population (Bayoumi *et al*., 1981; Holden and Mace, 1997; Tishkoff *et al*., 2007). Both −13907:G and −14009:G appear at their highest frequency in the Beja and it is possible that they emerged among these Sudanese populations. Another LP-associated SNP, −13915:G also appears at high frequency in the Beja population. The three alleles found in the Beja populations are on different haplotype backgrounds driving the frequency of putative lactose digesters to the highest seen in the area (Table 2). There is a clear extension of the haplotypes surrounding the derived alleles of the SNPs associated with LP (Figure 3). There is also an increase in LSBL-values close to the LP-associated region. Both of these signals suggest a selective event that drove the LP phenotype to a high frequency in the Beja population, that harbor several LP-associated alleles. The similar frequency of these three alleles in the Beni-Amer suggests that these SNPs are of similar age.

## Materials and Methods

A total of 221 individuals from 18 Sudanese and South Sudanese populations were selected for sequencing. These individuals have previously been investigated using microsatellites (Babiker *et al*., 2011) and dense SNPs (Hollfelder *et al*., 2017). A 316 base pair (bp) region of intron 13 of the *MCM6* gene was targeted for sequencing, encompassing all variants associated with LP (−13907, −13910, −13915, −14009, and −14010). Primer sequences were obtained from Coelho *et al*. (2009). DNA was extracted from Whatman FTA cards using Whatman protocol BD09 and BD01. PCR was performed using 0.625U AmpliTaq Gold DNA Polymerase, 1x Gold Buffer, 0.5mM dNTP mix, 2.5mM MgCl_2_, and 0.2µM of each primer per reaction in 30 cycles of 95*°*C at 15s, 55*°*C at 30s, and 72*°*C at 45s, with an initial deamination step of 10 minutes at 95*°*C and a final elongation of 5 minutes at 72*°*C. Sequencing was performed by the SNP&Seq Centre, SciLifeLab, Uppsala.

The obtained electropherograms were visually checked using GeneStudio and aligned to hg19 using MEGA7 (Kumar *et al*., 2016). Of the 221 individuals sequenced, 203 individuals gave successful sequencing results. All polymorphic sites were covered by concordant forward and reverse strands except for two individuals (one from each the Shaigia and the Bataheen populations) who had a successful result only with the forward-primer. All polymorphism peaks were unambiguous.

### Phasing and imputation to analyze haplotype structure

The sequencing results were added to 323,726 additional SNPs from chromosome 2, obtained from a filtered dataset of 3.9 million SNPs, typed on an Illumina HumanOmni5M Exome SNP array in a previous study (Hollfelder *et al*., 2017). This combined dataset was imputed and phased using fastPHASE version 1.4.0 (Scheet and Stephens, 2006). The number of haplotype clusters was set to 25, with 25 runs of the EM algorithm. The number of haplotypes sampled from the posterior distribution obtained from a particular random start of the EM algorithm was set to 100. We used the phase information to create a visualization of the haplotypes surrounding the LP control region (Figure 5). The R-package ‘rehh’ (Gautier and Vitalis, 2012) was used to create bifurcation plots visualizing the haplotype structure surrounding the LP-associated alleles.

### Locus specific branch length (LSBL)

Regions of extreme genetic differentiation on chromosome 2 were detected using LSBL (Shriver *et al*., 2004). LSBL estimates the branch length per locus by comparing pairwise *F*_*ST*_ values of three populations. This allows to detect in which of the three populations the genetic differentiation took place.

LSBL was calculated on the SASS populations from the Hollfelder *et al*. (2017) dataset as well as populations from the 1000 Genomes Project (1000 Genomes Project Consortium, 2015). The Hollfelder *et al*. (2017) dataset experienced a degree of allelic drop-out, which excludes the possibility of selection scans using haplotype based methods for this dataset. It was, however, shown that *F*_*ST*_ estimates on this diploid data set correlate strongly with a randomly haploidized version of the data set, therefore, measures such as LSBL can be used safely on the fully diploid data set (Hollfelder *et al*., 2017, SI).

We calculated Weir and Cockerham’s *F*_*ST*_ as implemented in plink v1.90 (Chang *et al*., 2015). LSBL was calculated for each locus on the SASS populations using two comparative non-LP populations (YRI and CHB), one African and one non-African to account for admixture in the SASS populations.

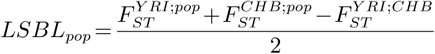

where *pop* is the test population. LSBL is calculated for each of the three combined populations. All SASS populations were tested, as well as MKK and CEU, which have been subjected to strong selection in the genomic region of the LP-associated alleles (Bersaglieri *et al*., 2004; Schlebusch *et al*., 2013). We computed the mean LSBL in non-overlapping 500 kilo base (kb) windows containing at least 50 SNPs and highlighted areas that are more than 3 standard deviations higher than the mean (Figure 4, S1, S2, S4, S5). A control was performed where negative *F*_*ST*_ estimates were exchanged to 0 (Hider *et al*., 2013). The treatment of negative *F*_*ST*_ estimates did not have an impact on the results (Table S1).

## Supplementary Material

Supplementary table S1 and figures S1–S10 are available at Molecular Biology and Evolution online (http://www.mbe.oxfordjournals.org/).

## Acknowledgments

We would like to thank the volunteering participants of this project that provided DNA samples. Sanger sequencing was performed at the Uppsala Genome Center, which is part of the Swedish National Genomics Infrastructure. The computations were performed on a high performance compute cluster at Uppsalas Multidisciplinary Center for Advanced Computational Science (UPPMAX). This work was supported by the Swedish research council (grant number 2018-05537 to MJ, 621-2014- 5211 to CS), the European Research council (ERC 759933 to CS), and the Knut and Alice Wallenberg foundation.

