## Supplemental Table and Figures for "The genetic variation of lactase persistence alleles in northeast Africa"

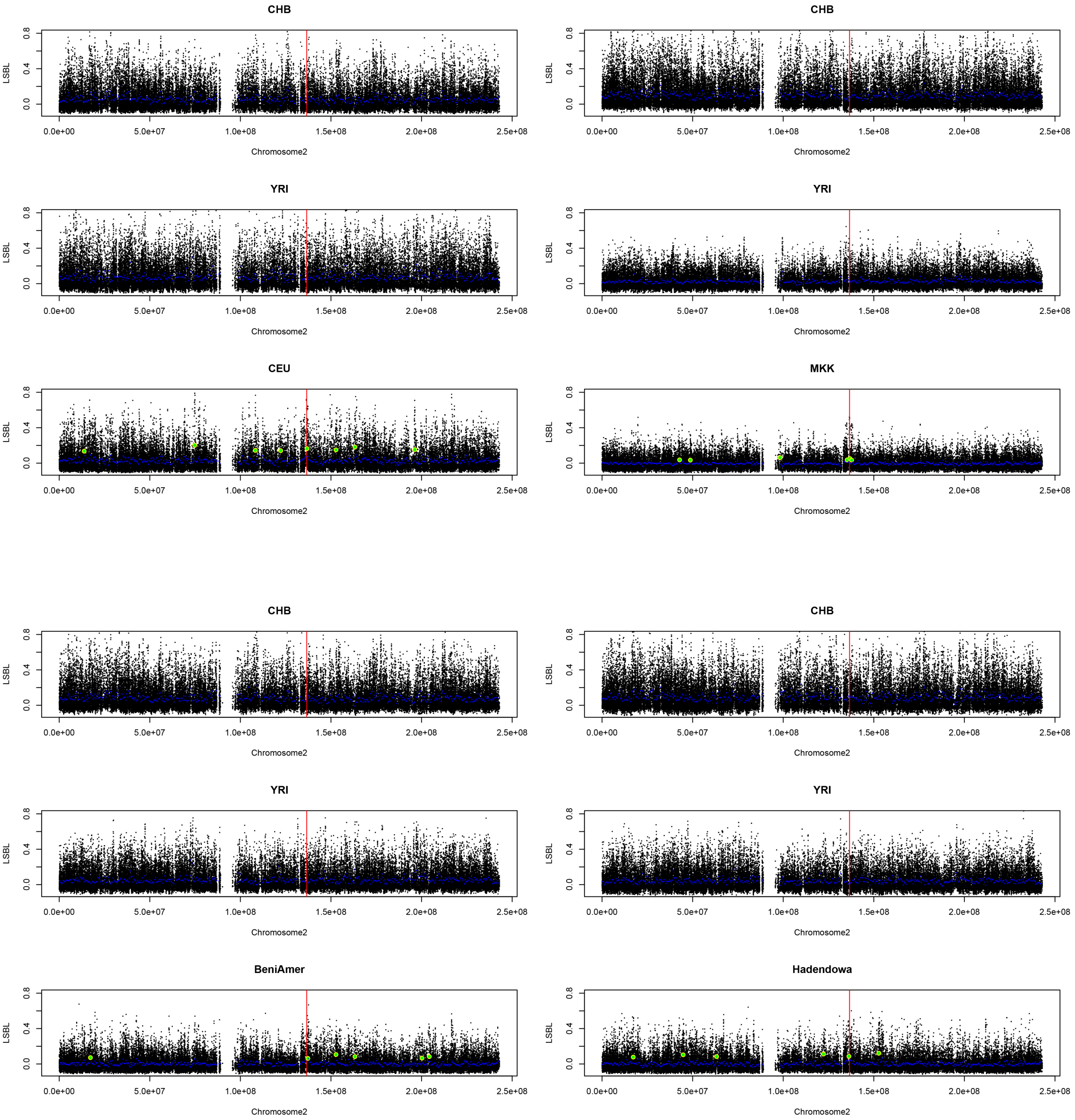


**FIG. 4.** LSBL result for CEU, MKK and the Beja populations. A group of three plots shows the population specific branch length for the combination CHB, YRI and X, where X is the population on the third plot of the group. The blue points indicate the means of 500kb windows, the larger green points show windows that deviate from the mean by more than 3 standard deviations. The red vertical line shows the position of -13910:C>T.

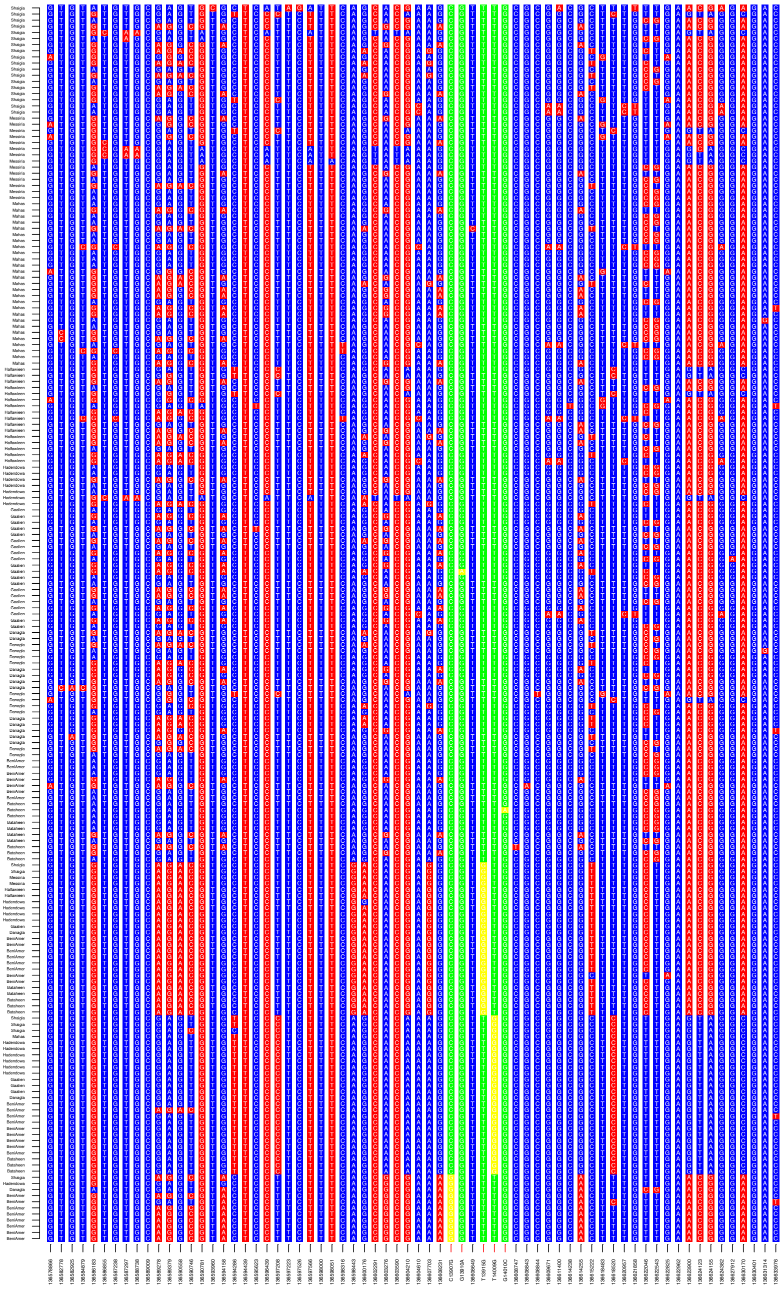

**FIG. 5.** Haplotype background. Only polymorphic SNPs are shown over an extension of 30kb in either direction of the LP-associated alleles in the populations that carry LP-associated alleles (Beja, Arab, Nubian). Each line represents one chromosome. Lactase-associated positions are highlighted in green/yellow, where yellow is the derived allele. The X-axis shows the bp-positions of the SNPs (except for the LP-associated SNPs) in GRCh37.p13.

### SUPPLEMENTAL FIGURES AND TABLES

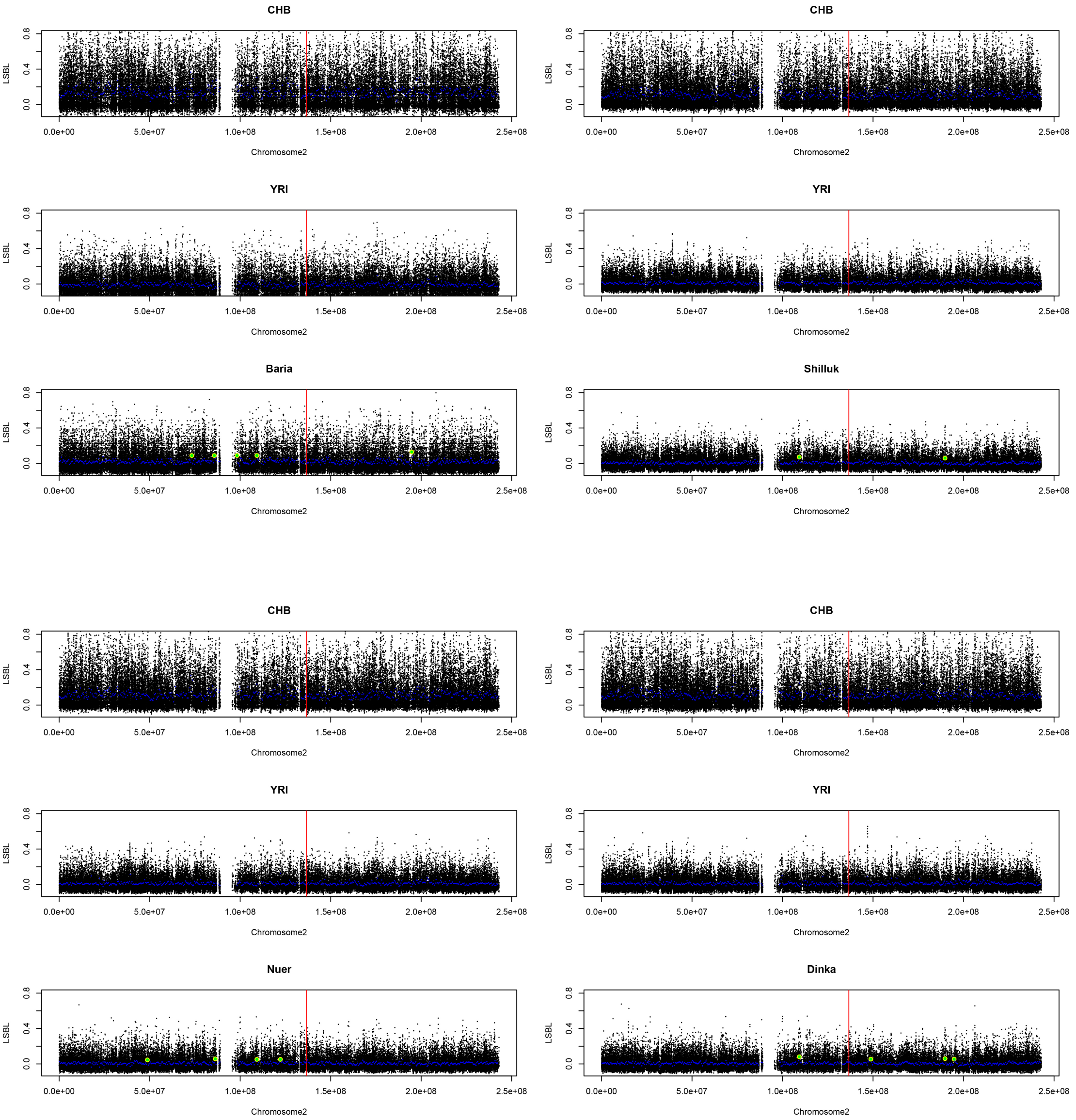


**FIG. S1.** LSBL of the Nilotes. A group of three plots shows the population specific branch length for the combination CHB, YRI and X, where X is the population on the third plot of the group. The blue points indicate the means of 500kb windows, the larger green points show windows that deviate from the mean by more than 3 standard deviations. The red vertical line shows the position of -13910:C>T.

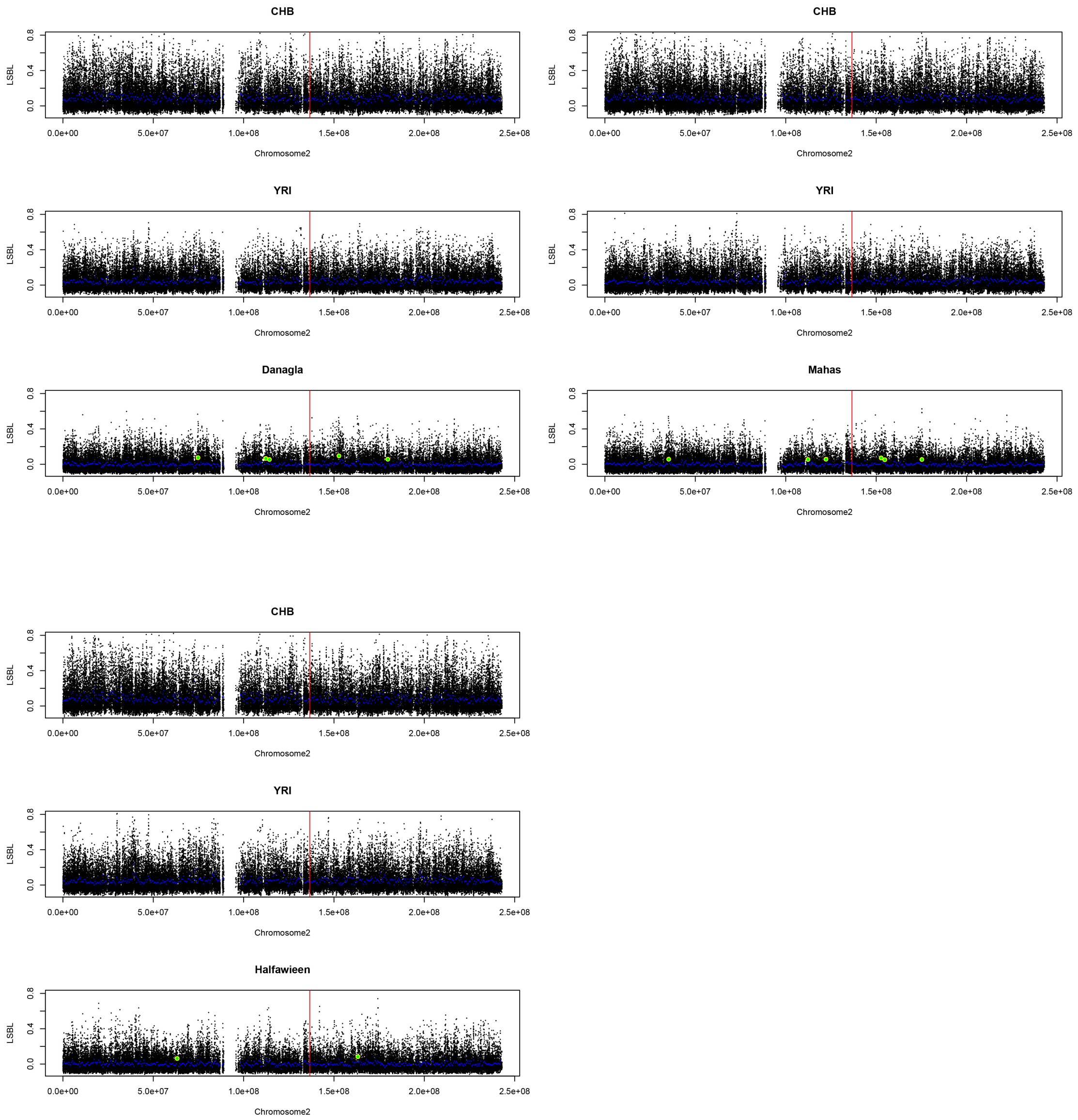


**FIG. S2.** LSBL of the Nubians. A group of three plots shows the population specific branch length for the combination CHB, YRI and X, where X is the population on the third plot of the group. The blue points indicate the means of 500kb windows, the larger green points show windows that deviate from the mean by more than 3 standard deviations. The red vertical line shows the position of 13910:C>T.

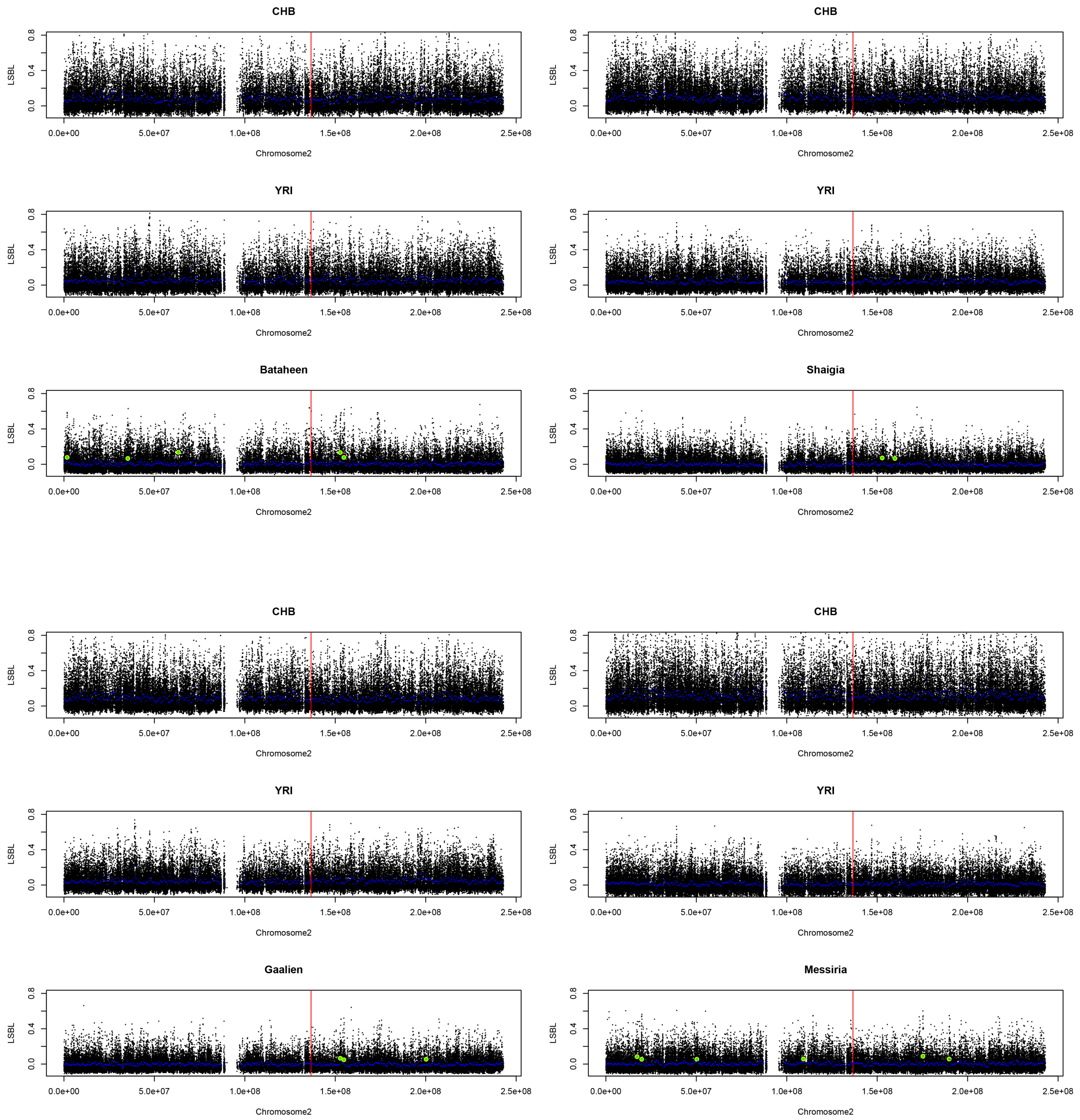


**FIG. S3.** LSBL of the Arab populations. A group of three plots shows the population specific branch length for the combination CHB, YRI and X, where X is the population on the third plot of the group. The blue points indicate the means of 500kb windows, the larger green points show windows that deviate from the mean by more than 3 standard deviations. The red vertical line shows the position of 13910:C>T.

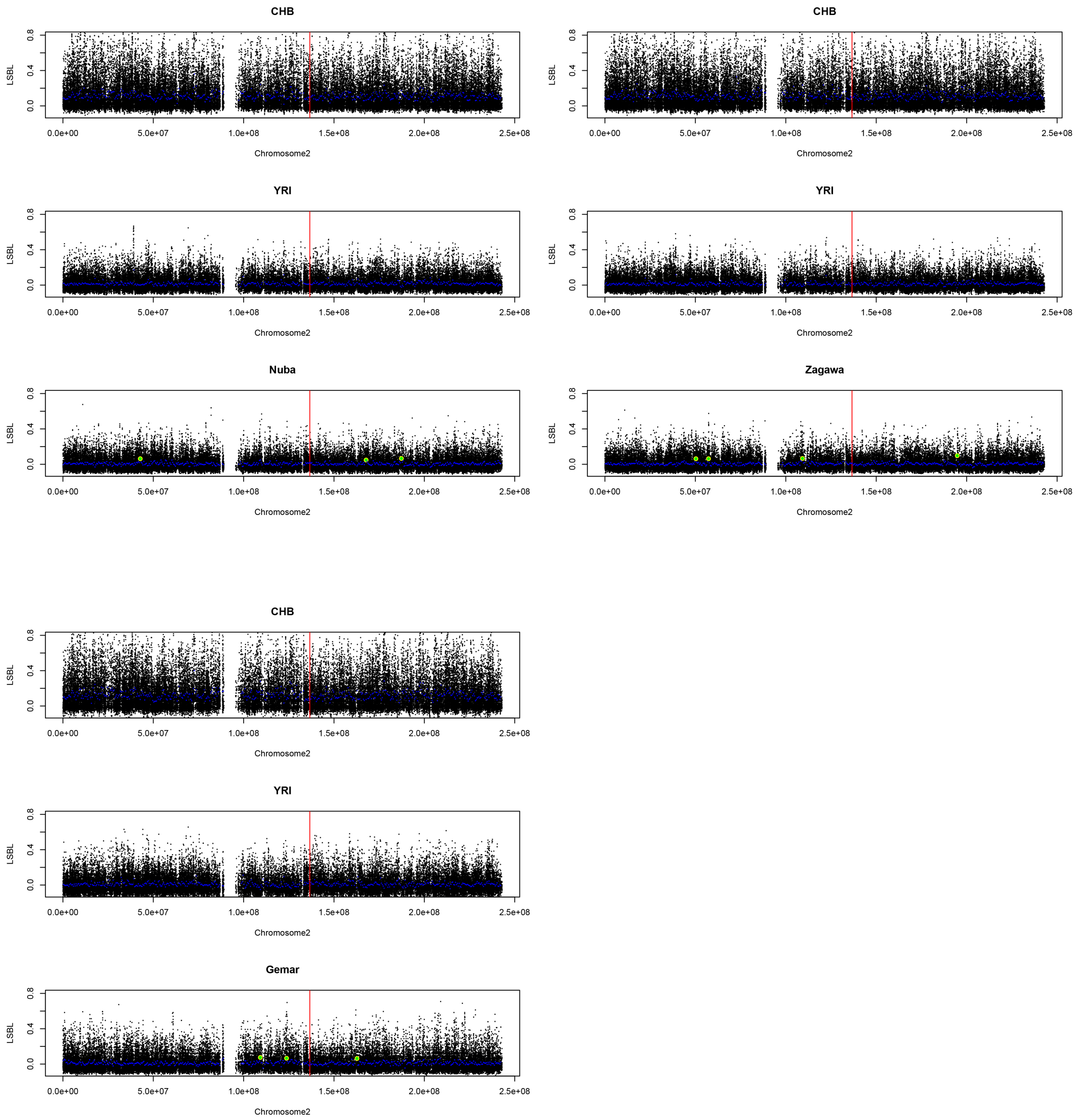


**FIG. S4.** LSBL of the Darfurian/Kordofonanian populations. A group of three plots shows the population specific branch length for the combination CHB, YRI and X, where X is the population on the third plot of the group. The blue points indicate the means of 500kb windows, the larger green points show windows that deviate from the mean by more than 3 standard deviations. The red vertical line shows the position of 13910:C>T.

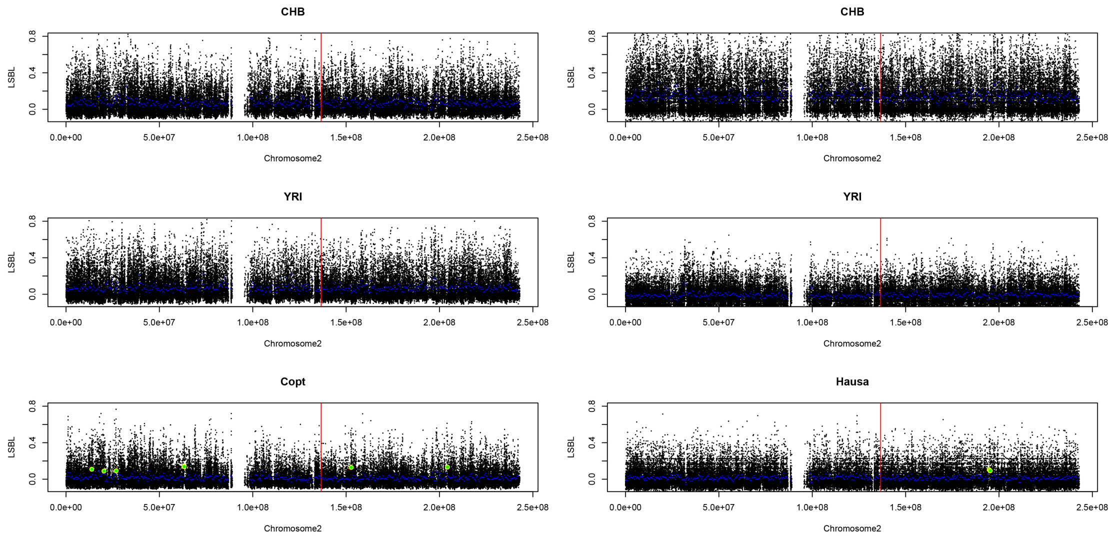


**FIG. S5.** LSBL of the migrant populations. A group of three plots shows the population specific branch length for the combination CHB, YRI and X, where X is the population on the third plot of the group. The blue points indicate the means of 500kb windows, the larger green points show windows that deviate from the mean by more than 3 standard deviations. The red vertical line shows the position of 13910:C>T.

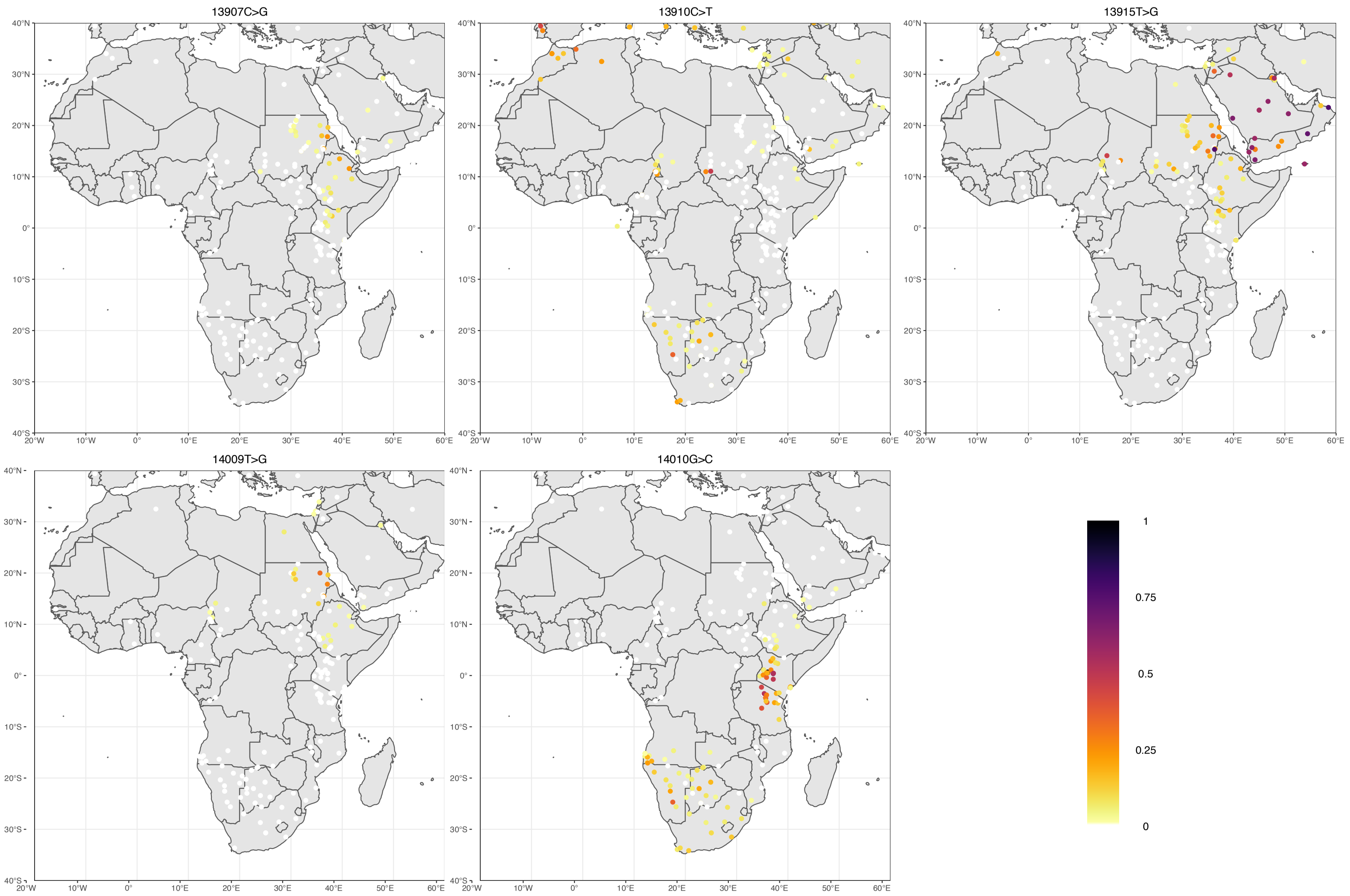


FIG. S6.: Allele frequency distribution of the derived alleles of the investigated SNPs in Africa. The white color signifies an allele frequency of zero.

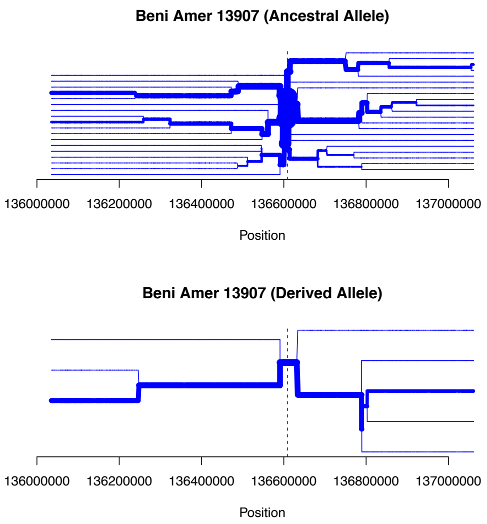


**FIG. S7.** Haplotype structure around -13907

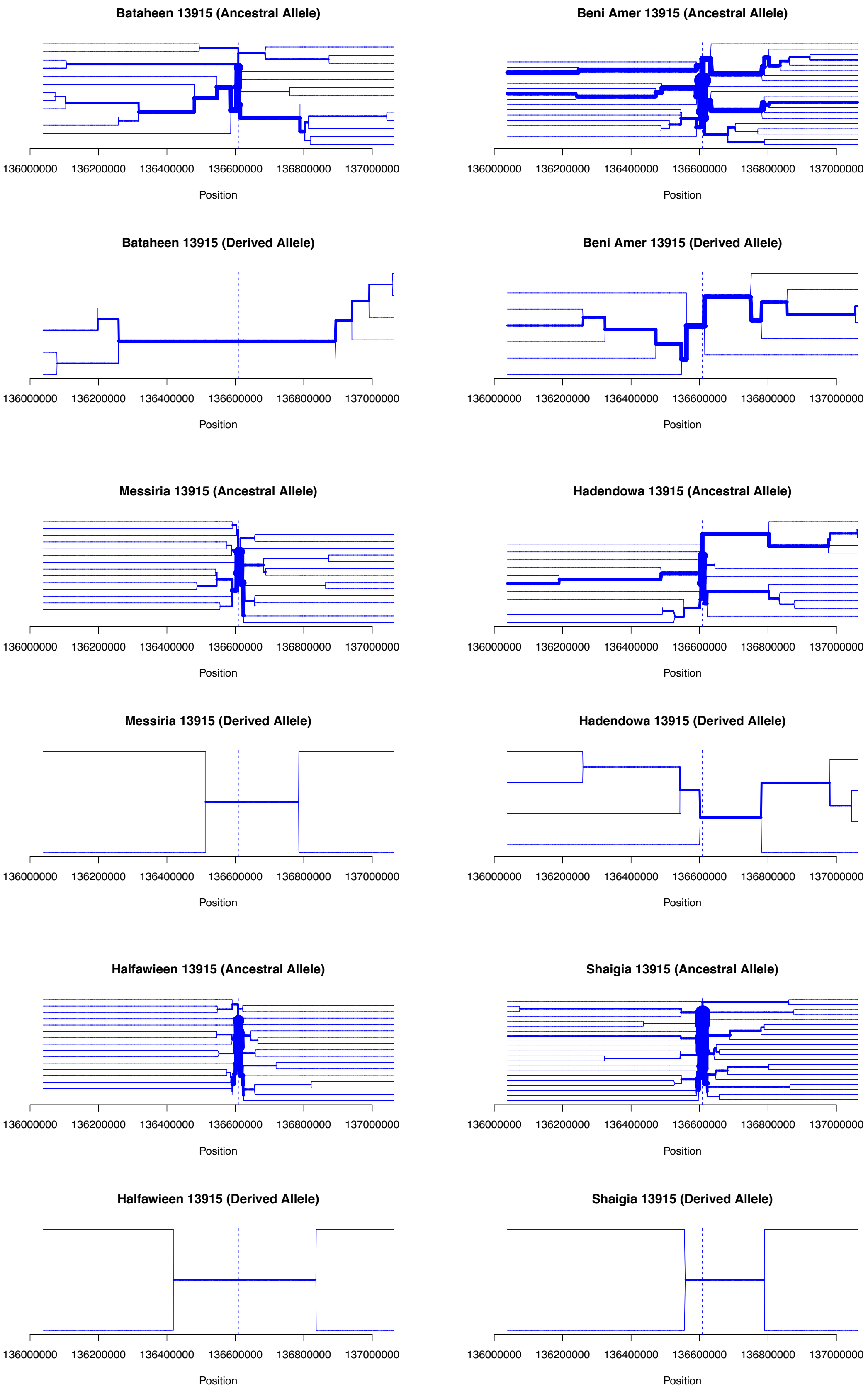


**FIG. S8.** Haplotype structure around -13915

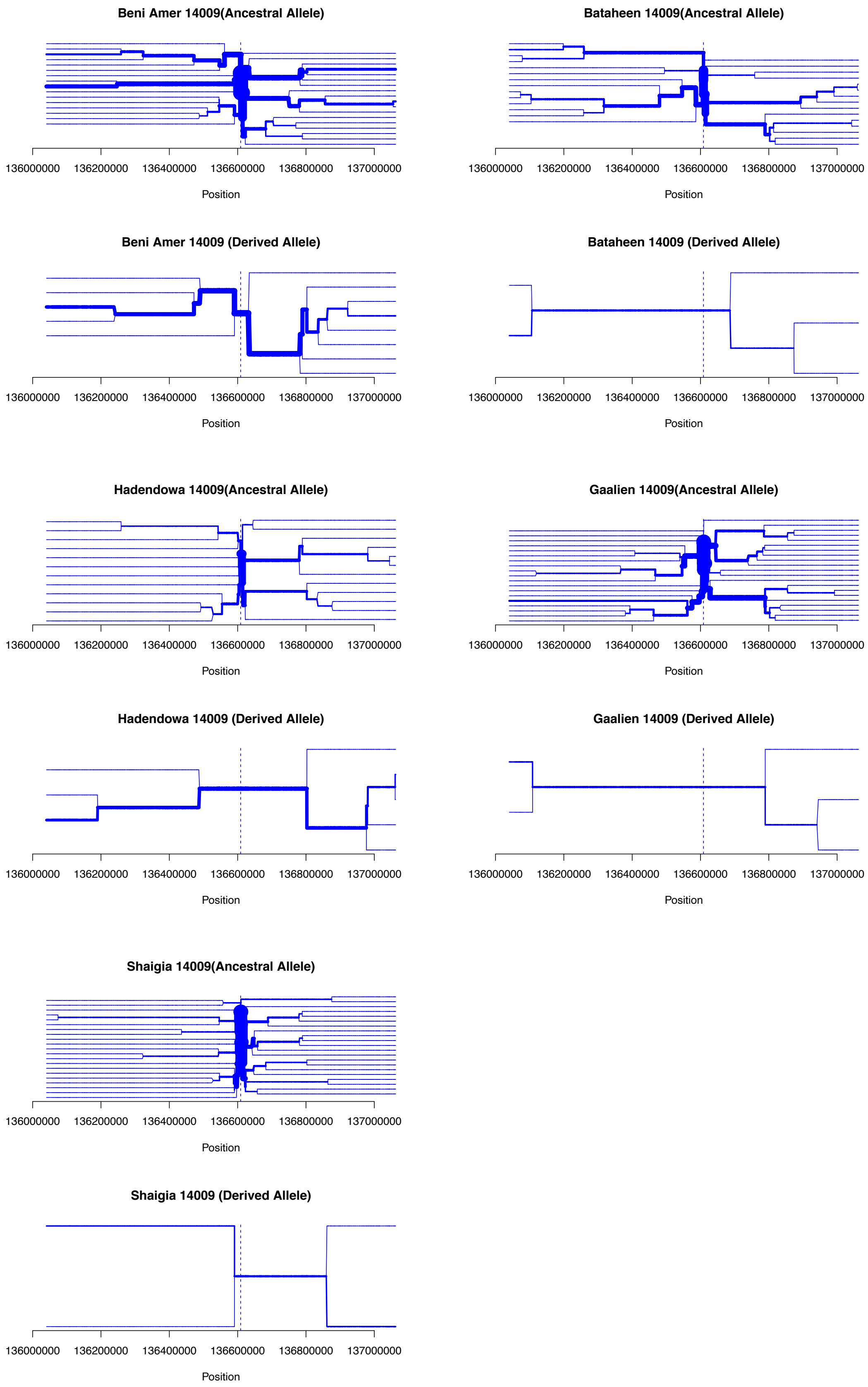


**FIG. S9.** Haplotype structure around -14009.

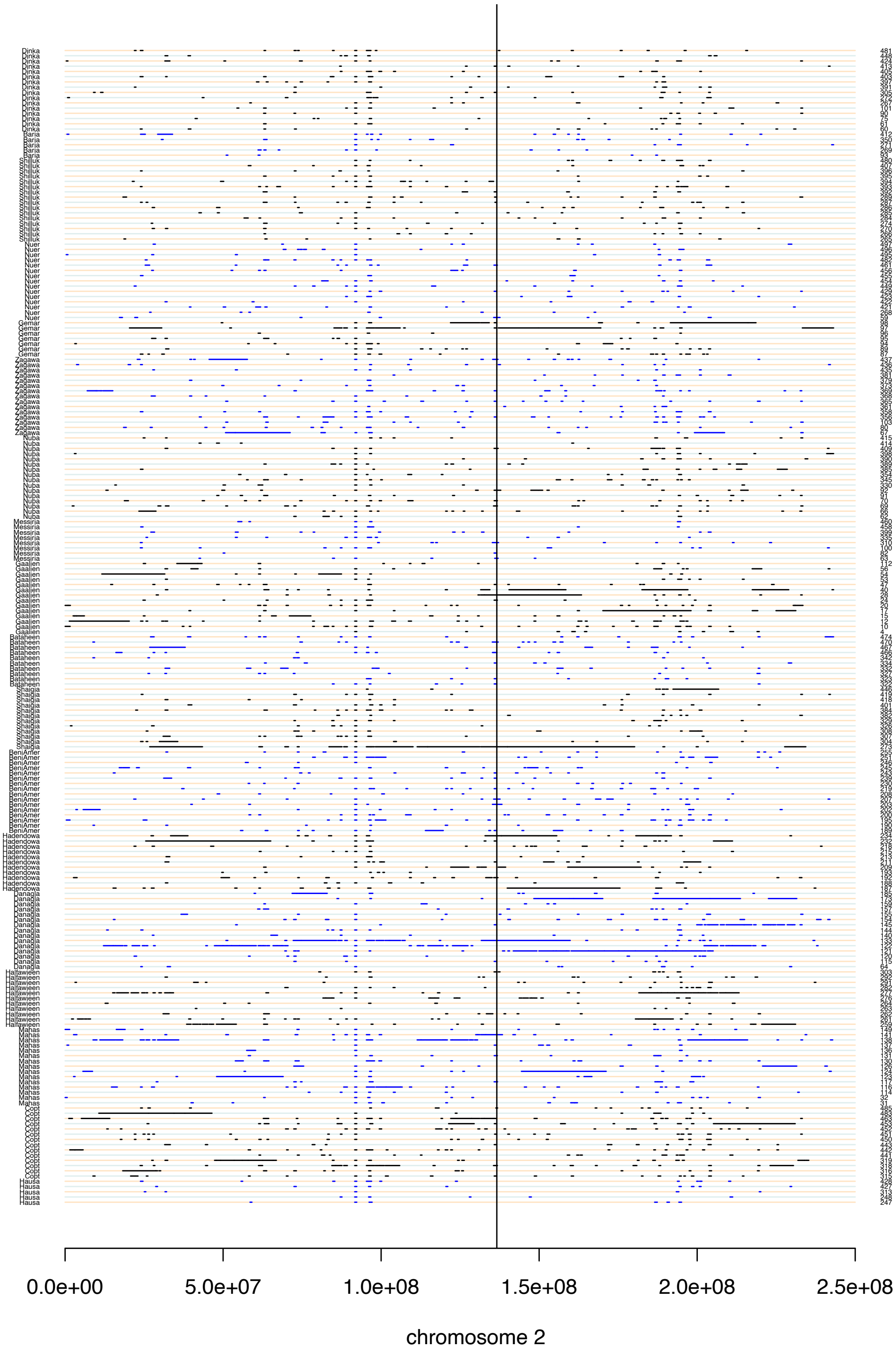


**FIG. S10.** Runs of Homozygosity on chromosome 2. The X-axis shows the chromosomal position, the vertical black bar shows the position of 13910:C>T. The Y-axis on the left shows the population, the right the individual ID.

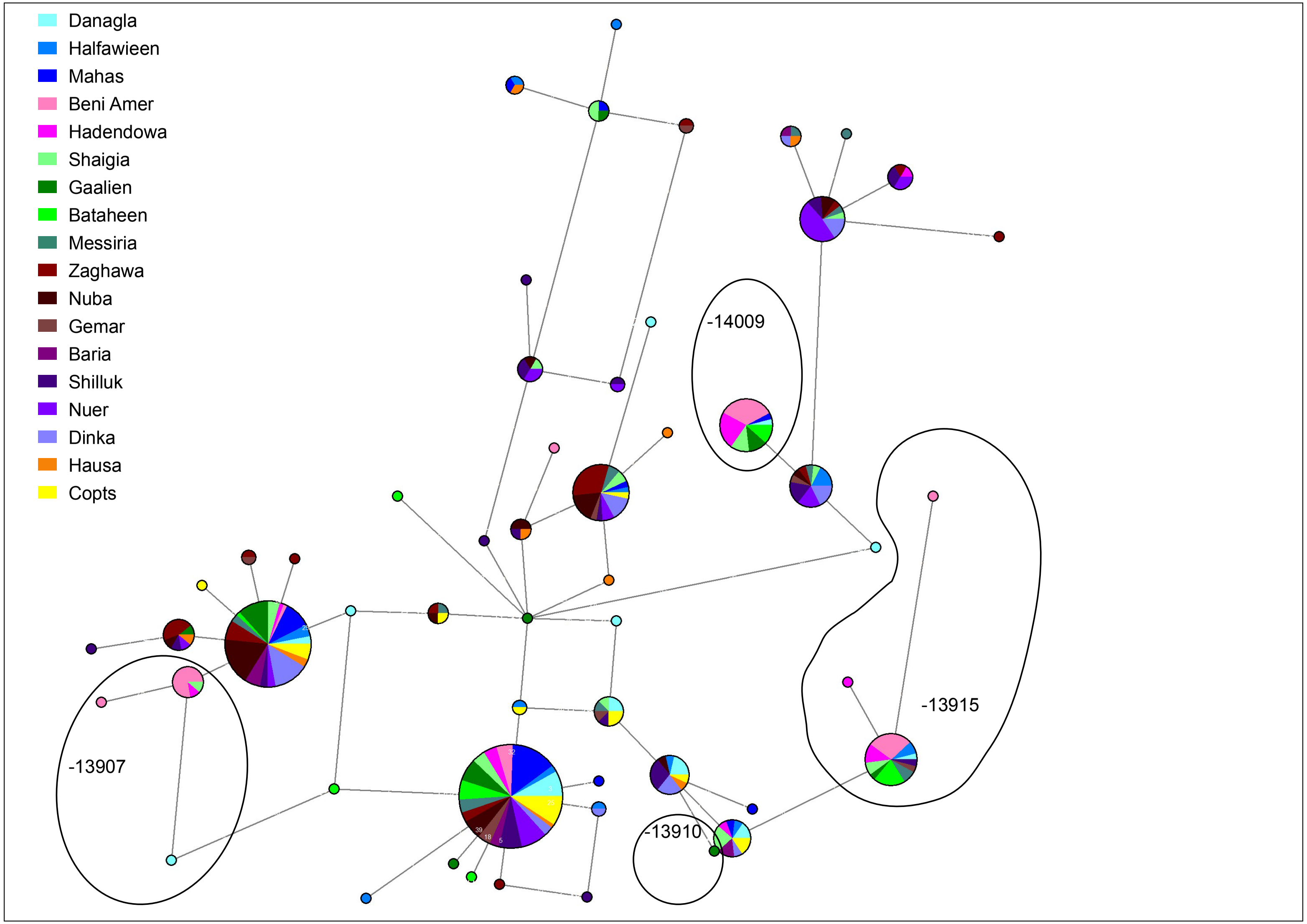


FIG. S11.: Haplotype network of polymorphic SNPs surrounding the Lactase SNPs, 10kb downstream and 20kb upstream.

**Table S1.** Regions that have significantly long LSBL. Region that includes the LP-associated alleles is marked red.

| 500 kb window | both $F_{ST}$ datasets | neg. $F_{ST}$ allowed | neg. $F_{ST}$ converted to 0 |
|---|---|---|---|
| 1518677-2018677 | Bataheen | | |
| 13518701-14018701 | CEU, Copt | | |
| 17018708-17518708 | Beni Amer, Hadendowa, Messiria | | |
| 19518713-20018713 | Messiria | | |
| 20018714-20518714 | Copt | | |
| 26518727-27018727 | Copt | | |
| 35018744-35518744 | Bataheen, Mahas | | |
| 42518759-43018759 | MKK, Nuba | | |
| 44518763-45018763 | Hadendowa | | |
| 48518771-49018771 | | Nuer, MKK | |
| 50018774-50518774 | Zaghawa | Messiria | |
| 57018788-57518788 | Zaghawa | | |
| 63018800-63518800 | Bataheen, Copt, Hadendowa, Halfawieen | | |
| 73018820-73518820 | Baria | | |
| 74518823-75018823 | CEU, Danagla | | |
| 84518843-85018843 | | | Shilluk |
| 85518845-86018845 | Baria | | |
| 86018846-86518846 | Nuer | | |
| 88518851-89018851 | | | Gemar |
| 98018870-98518870 | Baria, MKK | | |
| 108018890-108518890 | CEU | | |
| 109018892-109518892 | Dinka, Gemar, Messiria, Nuer, Shilluk, Zaghawa | Baria | |
| 112018898-112518898 | Danagla, Mahas | | |
| 114018902-114518902 | Danagla | | |
| 122018918-122518918 | CEU, Hadendowa, Mahas, Nuer | | |
| 123518921-124018921 | Gemar | | |
| 135018944-135518944 | MKK | | |
| 136018946-136518946 | Hadendowa | | |
| 136518947-137018947 | CEU, MKK | | |
| 137018948-137518948 | Beni Amer | | |
| 137518949-138018949 | MKK | | |
| 148518971-149018971 | Dinka | | |
| 152518979-153018979 | Bataheen, Beni Amer, CEU, Copt, Danagla, Gaalien, Hadendowa, Mahas, Shaigia | | |
| 154518983-155018983 | Bataheen, Gaalien, Mahas | | Hadendowa |
| 159518993-160018993 | Shaigia | | |
| 162518999-163018999 | Gemar | | |
| 163019000-163519000 | Beni Amer , CEU, Halfawieen | | |
| 167519009-168019009 | Nuba | | |
| 175019024-175519024 | Mahas, Messiria | | |
| 179519033-180019033 | Danagla | | |
| 187019048-187519048 | Nuba | | |
| 189519053-190019053 | Dinka, Messiria, Shilluk | | |
| 194519063-195019063 | Baria, Dinka, Hausa, Zaghawa | | |
| 195019064-195519064 | Hausa | | |
| 196019066-196519066 | CEU | | |
| 200019074-200519074 | Beni Amer, Gaalien | | |
| 204019082-204519082 | Beni Amer, Copt | | |

NOTE.—There are two regions that are distinguished in more than four populations. The first region (2:152518979-153018979) harbors the following genes: *NEB*, *ARL5A*, *CACNB4*, and *STAM2*. These Genes are associated with: nemaline myopathy 2, anxiety disorders, pancreatitis, waist circumference, lung neoplasms, altruism, body height, epilepsy, alopecia, hemoglobins, alcoholism, and chemo-therapy induced alopecia in breast cancer patients. The second region (2:109018892-109518892) contains the genes *GCC2* associated with child development disorders, *LIMS1*, *CCDC138*, and *RANBP2* which is associated with susceptibility to encephalopathy, and is downstream of *EDAR*, which is associated with hair and tooth morphology and ectodermal dysplasia.
